# Efficient workflow for validating homology-independent targeted integration-mediated gene insertion in rod photoreceptor cells to treat dominant-negative mutations causing retinitis pigmentosa

**DOI:** 10.1101/2023.11.08.566127

**Authors:** Akishi Onishi, Yuji Tsunekawa, Michiko Mandai, Aiko Ishimaru, Yoko Ohigashi, Junki Sho, Kazushi Yasuda, Keiichiro Suzuki, Juan Carlos Izpisua Belmonte, Fumio Matsuzaki, Masayo Takahashi

**Affiliations:** Laboratory of Retinal Regeneration, RIKEN Center for Biosystems Dynamics Research, Kobe Japan; Cell and Gene Therapy in Ophthalmology Laboratory, RIKEN Baton Zone Program, Kobe Japan; Laboratory for Cell Asymmetry, RIKEN Center for Biosystems Dynamics Research, Kobe Japan; Gene Expression Laboratory, Salk Institute for Biological Studies, La Jolla, USA; Division of Molecular and Medical Genetics, Center for Gene and Cell Therapy, The Institute of Medical Science, The University of Tokyo, Tokyo, Japan; Institute for Advanced Co-Creation Studies, Osaka University, Suita, Japan; Graduate School of Engineering Science, Osaka University, Toyonaka, Japan; Altos Labs, Inc., San Diego, California, USA; Kobe City Eye Hospital Research Center, Kobe, Japan; VCGT Inc., Kobe, Japan; Vision Care Inc., Kobe, Japan

## Abstract

Among the genome-editing methods for repairing disease-causing mutations resulting in dominant inhibition, homology-independent targeted integration (HITI)-mediated gene insertion of the normal form of the causative gene is useful because it allows the development of mutation-agnostic therapeutic products. For the rapid optimization and validation of highly effective HITI-treatment gene constructs against dominant-negative inheritance of inherited retinal dystrophy, we improved the gene constructs available in both plasmid and adeno-associated virus (AAV) vectors, and established a workflow that uses in vivo electroporation to verify the in vivo efficacy. By targeting the mouse Rhodopsin gene, we derived a construct in which HITI-mediated gene insertion occurs in 80%-90% of transduced mouse rod photoreceptor cells. This construct suppressed degeneration and induced visual restoration in the mutant mice. The HITI-treatment constructs for the rhodopsin gene were shown to be effective in AAV vectors, and this construction is available for the mouse Peripherin 2 gene. These findings suggest that the workflow reported here may be useful for the generation of HITI-treatment constructs for various target genes and for the development of gene therapy products.

## Introduction

Retinitis pigmentosa (RP) is the most prevalent form of inherited retinal dystrophy (IRD), and has a prevalence of 1 in 3,000–4,000 worldwide^1–4^. RP affects primarily the outer retinal layer and causes gradual degeneration of the photoreceptor cells^1,5,6^. Genetic diagnoses have uncovered more than 250 causative genes associated with IRD (see RetNet; http://sph.uth.tmc.edu/RetNet/ provided in the public domain by the University of Texas Houston Health Science Center, Houston, TX). These genes play pivotal roles in the function of photoreceptors and cells of the retinal pigment epithelium (RPE), and the mutant gene expression leads to impairment and degeneration^7^.

To treat patients with RP, several gene-targeted therapeutics have been proposed and tested^8–11^, depending on the inheritance patterns of disease-causing mutations^7^. Recessive mutations occur when the gene or protein expressed by the mutated gene allele is nonfunctional but does not influence the function of the wild-type (WT) gene allele. In individuals harboring a recessive mutation, gene therapy, also known as replacement therapy, is frequently used to introduce a normal gene from an external source^12^. On the other hand, dominant mutations constitute a type of inheritance (heterozygous) in which a mutation in one of the two alleles precipitates disease development.

Symptoms resulting from these mutations emerge in two primary ways: haploinsufficiency and dominant inhibition. Haploinsufficiency arises when the gene product expressed from one of the remaining alleles cannot sustain the original function, which thereby compromises the overall function because of reduced gene product levels. Dominant inhibition involves inhibition of the function of the WT gene or protein expressed by a mutated gene allele, which thereby disrupts normal cellular processes. Gene therapy for diseases caused by dominant mutations requires the supplementation of normal genes and repair of causal gene mutations^13^.

Genome-editing technology using gene-editing tools^14–16^ is used to repair genomic mutations that cause dominant inhibition and is therefore expected to provide fundamental treatment^17,18^. The homology-independent targeted integration (HITI) method is a gene-editing technique for repairing mutations by inserting or replacing a normal gene or editing the mutant base responsible for the mutation^19,20^. When a normal full-length sequence is introduced as a donor gene into a mutated target allele, the normal gene is expressed instead of the mutated gene. This method is considered to be “mutation agnostic” by enabling the same genome-editing target and donor cassette to treat all mutations, including novel mutations^21,22^. However, few studies have reported on the development of the currently available highly efficient HITI-treatment constructs used for target gene insertion into a donor cassette to stop the mutant gene expression and that are expressed at the same level as the normal allele and the treated photoreceptors.

Here, we report an efficient method for validating gene constructs for HITI-mediated exogenous gene insertion. We focused on targeting the rhodopsin (*RHO*) gene, the leading cause of autosomal dominant retinitis pigmentosa (AdRP)^23,24^. Using *Streptococcus pyogenes* Cas9 (SpCas9) as a gene-editing tool and in vivo electroporation as a gene-transfer technique, we identified SpCas9 guide RNA (gRNA) target sequences with high insertion efficiency in vivo, although the efficiency did not correlate fully with the scores obtained from in silico gRNA finders. The inserted normal genes from the donor vectors expressed with chimeric introns and the 3′-UTR (untranslated region) of the target genes showed therapeutic efficacy in *Rho*-mutant mice. Given that the HITI-treatment constructs targeting the mouse peripherin2 (*Prph2*) gene^25^, produced using the same methodology as that for *Rho*, were shown to be effective, this workflow for constructing HITI-mediated gene insertions may be applied broadly in gene-targeted therapeutics for RP.

## Results

### Design and validation of the gene constructs for highly efficient HITI-mediated insertion

Many genes responsible for RP are enriched in rod photoreceptor cells^13^. Therefore, we attempted to establish an experimental procedure for the rapid construction and validation of a highly efficient HITI-mediated insertion of exogenous genes into rod photoreceptor cells. Because gene mutations are often contained in the first exon of coding sequences (CDS), it is desirable to insert a donor gene cassette containing the normal genes and polyadenylation (poly(A)) sequences into the 5′-UTR region. More than 100 mutations have been reported in exons 1–5 in of the human *RHO* locus^24^.

First, we selected SpCas9 gRNA target sequences to introduce double-stranded DNA breaks (DSBs) into the 50 bp 5′-UTR region proximal to the *Rho* start codon, which is evolutionarily nonconserved (Fig. 1A). Using grID (gRNA Identification) database, in silico SpCas9 gRNA finders^26^ (http://crispr.technology), we selected the three highest-scoring gRNAs with grID scores ranging from 600 to 800 (Supplemental Table S1). To evaluate the cleavage efficiency of the three target sequences in vitro, we performed a single-strand annealing (SSA) assay, in which DSB efficiency was validated using SSA-mediated *EGFP* reconstitution of pCAG-EGxxFP plasmids carrying 5′- and 3′-*EGFP* fragments with the three gRNA target sequences (Fig. 1B). The cleavage efficiencies of gRNA1 and gRNA3 were similar, whereas gRNA2 exhibited low EGFP expression.

**Figure 1.**
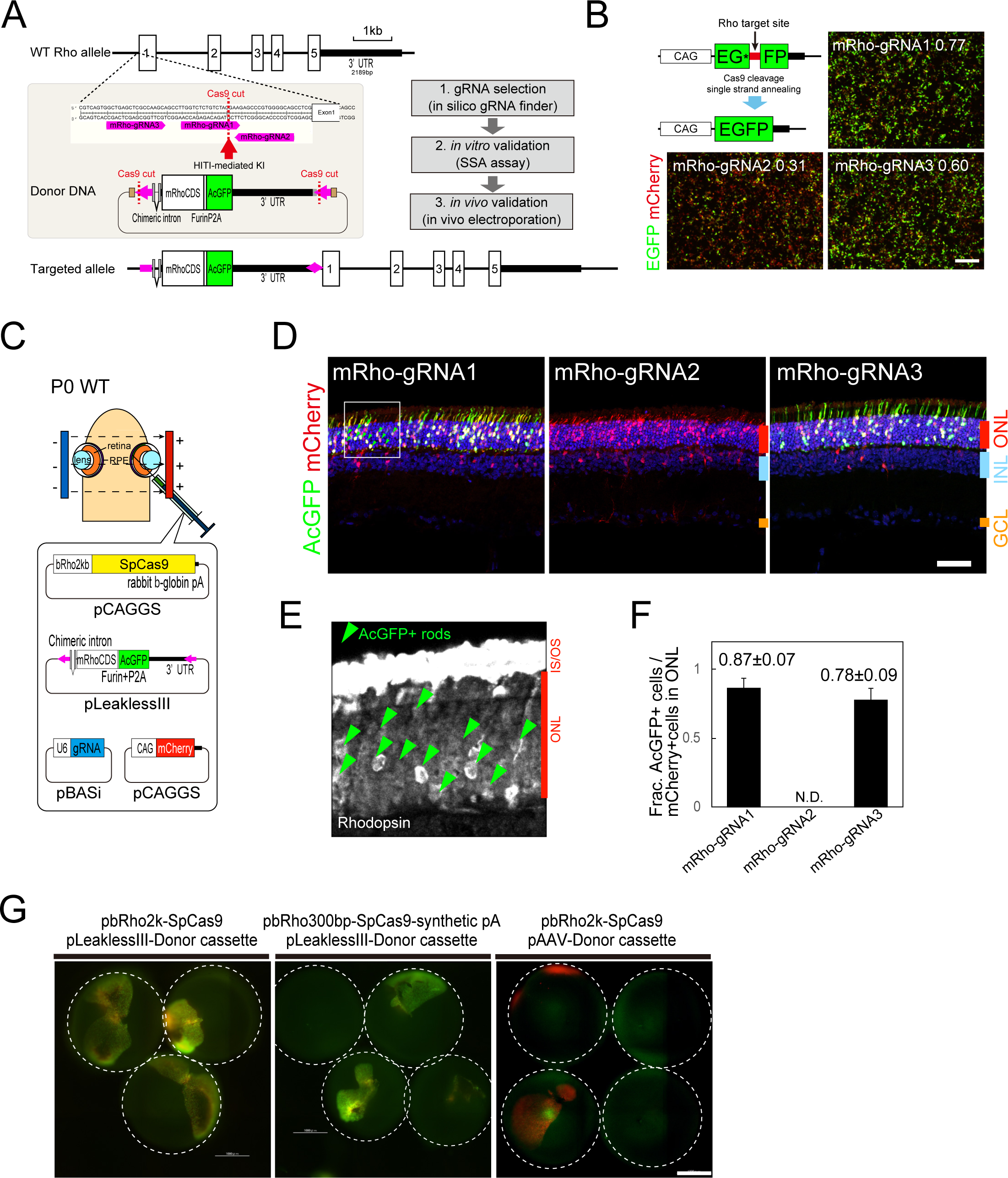
Optimization and validation of highly efficient HITI-treatment gene constructs targeting the mouse *Rho* locus. (A) Schematic illustration and workflow of the Cas9-driven HITI-treatment gene construction. Each donor cassette containing one of the three gRNA sequences (pink) selected from the 5′-UTR sequence of the mouse *Rho* locus at both ends was prepared. The targeted allele represents the gene structure when the HITI-mediated gene insertion occurs. (B) SSA assay for in vitro evaluation of the cleavage efficiency of the three gRNA sequences targeting Rho. The illustration indicates the SSA-mediated reconstruction of pCAG-EGxxFP fragments by Cas9 cleavage, and the photographs represent HEK293T cells 48 h after transfection with pCAG-EGxxFP, pCAG-mCherry, pCAG-SpCas9, and the three gRNA plasmids. Numbers represent EGFP:mCherry intensity ratios measured by ImageJ. Scale bar: 500 μm. (C) Schematic illustration of HITI-mediated donor insertion by in vivo electroporation into the mouse retina. The construction of each plasmid with a vector backbone is shown. The plasmid cocktail was injected into the subretinal space of postnatal day 0 (P0) mouse retinas. (D) Immunohistochemistry sections of P21 mouse retinas electroporated in vivo at P0 with three plasmid cocktails targeting different mRho-gRNA1 (left), mRho-gRNA2 (middle), and mRho-gRNA3 (right) sequences. Electroporated cells are mCherry positive, and most cells in the outer nuclear layer (ONL) express AcGFP in retinal sections targeting mRho-gRNA1 and mRho-gRNA3 sequences. INL, inner nuclear layer; GCL, ganglion cell layer. Scale bar, 50 μm. (E) High-power image of section immunohistochemistry with anti-rhodopsin antibody (white) of the square area shown in Fig. 1D. Arrowheads indicate AcGFP-expressing cells. (F) Fraction (Frac.) of AcGFP-positive cells in mCherry-positive ONL cells from three different retinal tissue sections. Numbers, the mean ± SD (n = 6); N.D., not detected. (G) P21 mouse eyecups electroporated in vivo with plasmid cocktails targeting the gRNA sequence, in which the bRho2k promoter was shortened (bRho300bp) to express Cas9 (middle), and the vector backbone for the donor cassette was switched to pAAV (right). The dashed circles represent each dissected eyecup. Scale bar, 1 mm.

Next, donor plasmids were designed. Rhodopsin accounts for 95% of the proteins constituting the outer segments^27^, and it is important to maintain high expression levels. Previously, the use of chimeric introns^28–30^ was effective in increasing exogenous gene expression at the *Rho* locus in knock-in (KI) mice ^31,32^. A poly(A) sequence derived from the mouse *Rho* locus was also used. Analysis of mouse *Rho* mRNA revealed the presence of multiple poly(A) signals in the 3′-UTR, which suggested the stability of and increase in *Rho* mRNA transcription^33^. The reverse sequence of each gRNA sequence was added to both ends of the gene cassette and inserted into pLeaklessIII, a donor plasmid vector developed to prevent leaky transcription from the plasmid backbone^34,35^.

### HITI-mediated gene insertion observed in 80%–90% of electroporated rod photoreceptors

We performed in vivo electroporation to rapidly evaluate the in vivo insertion efficiency of HITI-mediated gene insertion in mouse rod photoreceptor cells. We used pRho2k-SpCas9, in which SpCas9 was expressed from a 2 kb promoter of bovine rhodopsin^36^ in differentiated mouse rod photoreceptor cells (Fig. 1C). Plasmid vectors for gRNA expression (U6-gRNA1, 2, and 3), donor constructs, and reporter expression (pCAG-mCherry) were coelectroporated in the postnatal day 0 (P0) newborn mouse retina, in which the plasmid DNAs can be transferred into the nucleus because the retinal cells of the future photoreceptor layer (outer neuroblastic layer) are in an undifferentiated dividing state. After these cells differentiate into rod photoreceptor cells in a nondividing state, SpCas9 is expressed from the rhodopsin promoter.

In the P21 eyecups, *Aequorea coerulescens* GFP (AcGFP) expression was observed in gRNA1- and gRNA3-treated cells (Supplemental Fig. S1), which indicated HITI-mediated KI of the donor cassettes in the correct direction. Section immunohistochemistry (IHC) of these retinas showed that AcGFP-positive cells were localized at the photoreceptor layer, whereas mCherry signals were observed in the inner nuclear layer where other retinal neurons were developing (Fig. 1D). AcGFP-expressing cells were also coimmunostained with anti-rhodopsin antibodies, and the IHC images showed that mRhoCDS and AcGFP in the donor plasmids were inserted and expressed from intrinsic *Rho* promoters in the genome (Fig. 1E). By contrast, the eyecups electroporated with gRNA2 were mCherry positive, but AcGFP fluorescence was not detectable (Fig. 1C, Supplemental Fig. S1), which suggested that gRNA2 was ineffective in vivo. The efficiency of each HITI-mediated donor insertion was estimated from the percentage of AcGFP-positive cells to the number of mCherry signals localized in the photoreceptor layer (Fig. 1F). The percentage was 80%–90%, which suggested that HITI-mediated gene insertion occurred in most electroporated photoreceptor cells.

Given that the Rho2k-Cas9 cassette is about 7 kb in length, which is greater than the AAV packaging limit, we examined HITI efficiency using a 300 bp rhodopsin proximal promoter. The AcGFP fluorescence of P21-electroporated retinal eyecups was similar to that of the Rho2k promoter (Fig. 1G), which indicated that the 300 bp promoter was available. However, when the plasmid backbone of the donor vector was switched to pAAV instead of pLeaklessIII, AcGFP fluorescence was not detected. This result indicated that the type of plasmid backbone influenced HITI efficiency.

### Monoallelic HITI-mediated insertion of rod photoreceptors

HITI-mediated gene insertion occurs in either a monoallelic or a biallelic manner in each rod photoreceptor. To compare the percentages of monoallelic and biallelic insertions in electroporated WT mouse rod photoreceptors, we performed single-cell genotyping using dissociated HITI-treated rod photoreceptors. For this experiment, we added the nuclear localization signals (NLSs) to AcGFP and mCherry (Fig. 2A) because AcGFP and mCherry localized in dissociated rod outer segments increased the error in cell sorting. At P56, we collected 48 cells from the AcGFP- and mCherry-double-positive fractions directly into 96-well plates and amplified the WT and KI fragments using primers specific to both sequences (Fig. 2B). Next, we performed nested PCR using primers to amplify either the WT or KI fragments (Fig. 2B). The cell samples in which only the KI band was amplified showed biallelic insertion, whereas those in which both the WT and KI bands were amplified were monoallelic (Fig. 2C, Supplemental Fig. S2).

**Figure 2.**
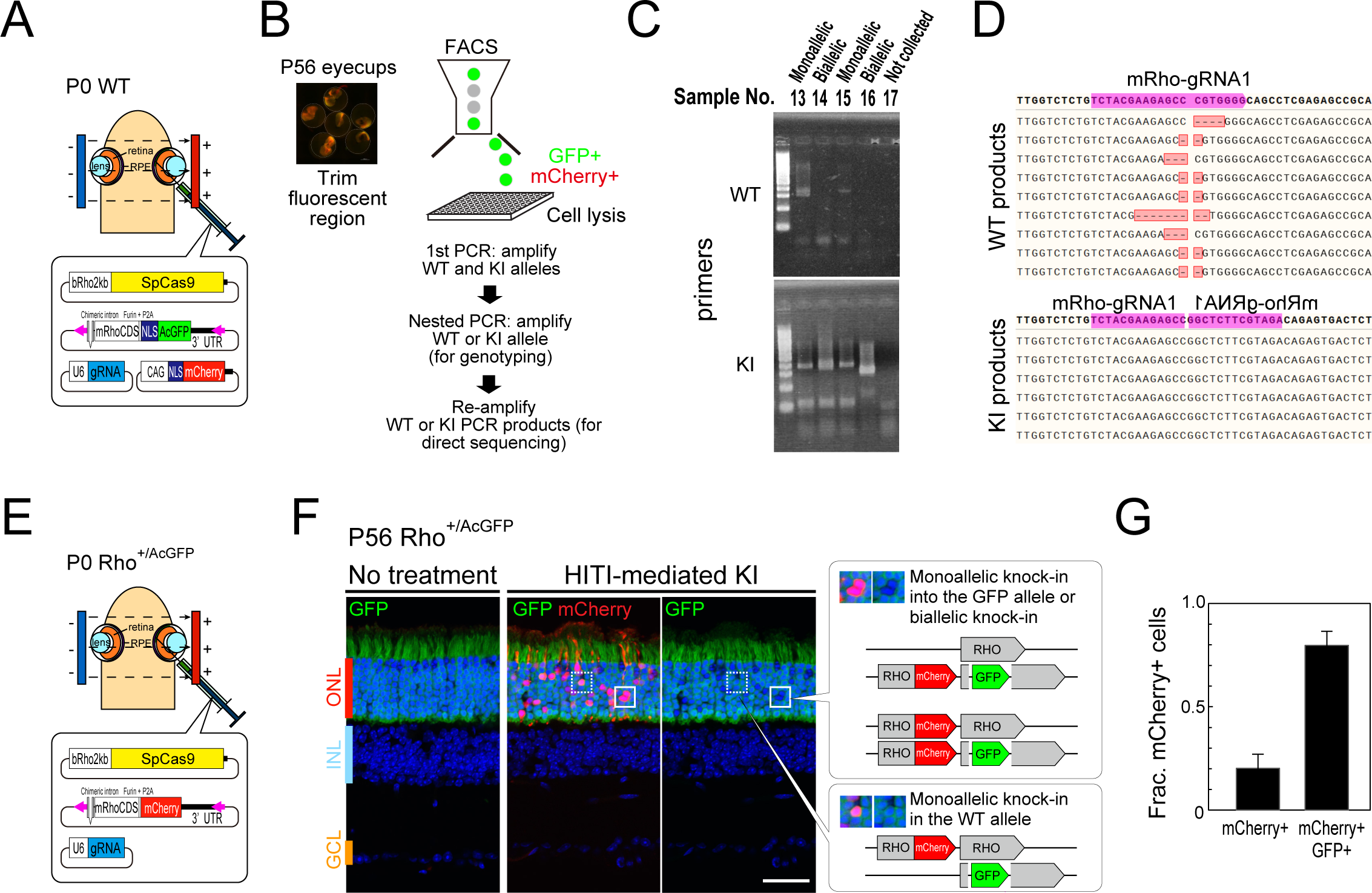
Validation of monoallelic and biallelic insertions in HITI-treated rod photoreceptor cells. (A) Schematic illustration of HITI-treatment gene construction for cell sorting. Nuclear localization signals (NLS) were added to the fluorescent markers in the donor and reporter plasmids. (B) Schematic illustration of single-cell sorting and PCR genotyping. The fluorescent region of P56-electroporated retinas was trimmed and dissociated, and 48 cells were sorted using FACS from the GFP- and mCherry-double-positive fractions. WT and KI alleles were amplified from the first PCR, and each allele was amplified using nested PCR. (C) Electrophoresis gel images of the nested PCR of WT and KI alleles of sample numbers 13–17 of the 48 samples tested. (D) Direct sequencing of the reamplified WT (upper) and KI (lower) PCR products. The sequences around the mRho-gRNA1 target (purple shading) are shown. (E) Schematic illustration of the de novo validation of the donor vector insertion. (F) P56-electroporated Rho^+/AcGFP^ retinal sections. AcGFP-positive and -negative cells were observed in HITI-treated mCherry-positive photoreceptor cells. Scale bar: 50 μm. (G) Fraction (Frac.) of mCherry- and/or AcGFP-positive cells counted in retinal sections. Data represents means□±□SD (n = 4).

We detected longer and shorter PCR fragments in some samples, possibly because of intermolecular annealing of the uncleaved donor plasmids (Fig. 2C). We then reamplified the PCR fragments and sequenced them (Fig. 2D). Using direct sequencing of the WT PCR fragments, we detected deletions in the gRNA1 cleavage region. Using direct sequencing of the KI PCR fragments, we detected nonhomologous end joining (NHEJ) between the *Rho* genomic target and donor cassettes. The electropherogram of each sample was uncontaminated, which indicated that the PCR fragments were amplified from a single allele.

Next, to verify the allele-specific HITI-mediated KI in vivo, we performed HITI insertion into Rho^+/AcGFP^ heterozygous mice in which *AcGFP* was inserted into intron 2 at the *Rho* locus (Supplemental Fig. S3). Because the KI mice expressed AcGFP in all rod photoreceptor cells, we performed HITI-mediated KI using donor cassettes containing mCherry (Fig. 2E). In the P56 electroporated Rho^+/AcGFP^ retinas, we observed that HITI-treated mCherry-positive cells in the outer nuclear layer (ONL) and that nonelectroporated cells maintained AcGFP expression (Fig. 2F). Some mCherry-positive cells lost AcGFP expression, which indicated insertion of a biallelic donor cassette to both alleles or monoallelic donor cassette insertion into the *AcGFP* knock-in allele. The presence of mCherry-positive cells that coexpressed AcGFP indicated the insertion of a monoallelic donor cassette into the WT allele (Fig. 2F). The percentage of mCherry- and AcGFP-double-positive cells was about 80%, which suggested that HITI-mediated KI was more monoallelic.

### Normal expression of *Rho* and suppressed rod photoreceptor degeneration of HITI-treated rod photoreceptors in *Rho* mutant mice

We confirmed the coexpression of mRhoCDS and AcGFP in electroporated WT mouse retinas. We then performed HITI-mediated donor cassette insertion into the Rho^P23H/P23H^homozygote mutant mouse retina to examine whether mRhoCDS expression suppresses rod photoreceptor cell degeneration (Fig. 3A). Because AdRP mutations cause photoreceptor degeneration, even in heterozygotes, we used P23H homozygous mice to quickly observe the suppressive effect against rod photoreceptor degeneration^37,38^ and to confirm the expression and localization of the inserted normal *Rho* gene.

**Figure 3.**
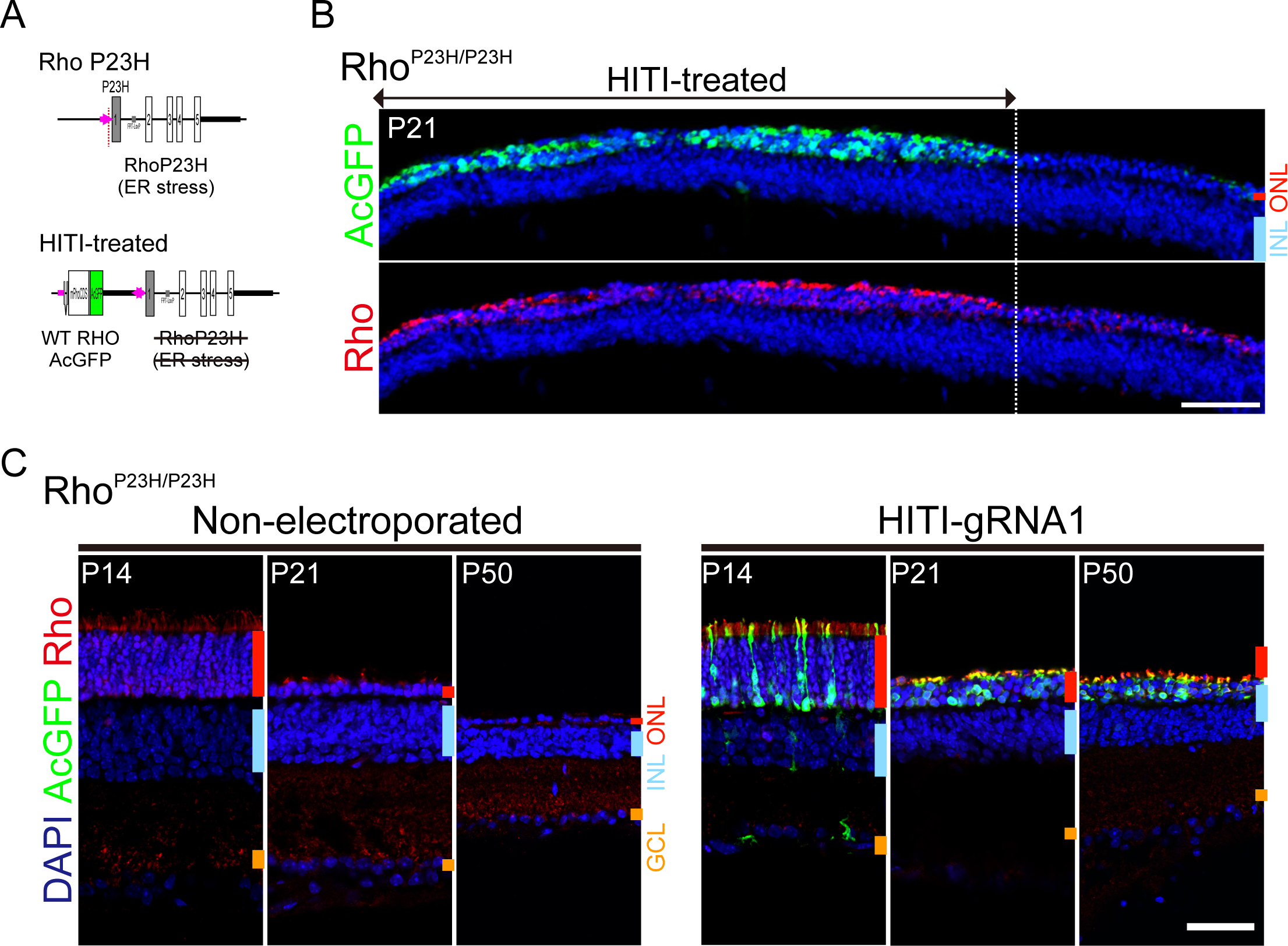
HITI-mediated normal rhodopsin knock-in suppression of rod photoreceptor degeneration in Rho^P23H/P23H^ mice. (A) Schematic illustration of HITI-mediated gene repair of the mouse Rho^P23H^ locus. (B) Low power image of section immunohistochemistry (IHC) of HITI constructs electroporated in vivo into Rho^P23H/P23H^ mice. Immunostaining of AcGFP (upper) and rhodopsin (lower) antibodies were shown. Scale bar: 50 μm. (C) Section IHC of P14, P21, and P50 retinas from P23H mice coimmunostained with AcGFP (green) and rhodopsin (red) andibodies. Scale bar: 50 μm.

The P21 Rho^P23H/P23H^ mouse retinas electroporated in vivo with HITI plasmids at P0 (shown in Fig. 1B) are shown in Fig. 3A. The AcGFP-positive ONL was thicker than the surrounding ONL, where AcGFP was negative, which suggested that electroporated rod photoreceptors were protected against degeneration. The electroporated rods were coimmunostained with anti-AcGFP and -RHO antibodies. RHO protein was preferentially localized outside the ONL, whereas RHOP23H mutant proteins were mislocalized. IHC sections obtained at several developmental time points showed that HITI-treated rod photoreceptors were maintained at P21 and P50, but nonelectroporated retinas showed progressive photoreceptor degeneration (Fig. 3C).

### Recapitulation of HITI-mediated KI into the mouse Rho locus using AAV vectors

Because electroporation has difficulty in transferring genes directly into cell nuclei in the nondividing state, AAV is often used as a vector for in vivo gene therapeutic agents. We examined whether the HITI-treatment constructs that worked in the electroporation-based validation could be switched into the AAV vectors (Fig. 1). We used the type 8 AAV serotype transduced into adult mouse photoreceptor cells. First, we packaged the HITI gene constructs into two AAV8 vectors, in which the donor and U6-gRNA cassettes were integrated into a single AAV8, according to a previous HITI study^20^ (Supplemental Fig. S5). AAV8 cocktails were injected into 2-month-old C57BL/6 mice. No AcGFP signals were observed in the eyecups after 1 month (Supplemental Fig. S5).

We next packaged the donor gene and gRNA expression cassette into separate AAV8 capsids to construct three AAV8 capsids (Fig. 4A). Because the U6-gRNA cassettes are about 0.4 kb, which is small compared with the AAV package size, two U6-gRNA cassettes were placed side by side across the woodchuck hepatitis virus posttranscriptional regulatory element (WPRE) and packaged into self-complementary AAV (scAAV). We inserted WPRE because AAV plasmids without the WPRE sequence could not be obtained in sufficient quantities using Midi/Maxiprep plasmid purification. The three AAV8 vectors were injected subretinally into 2-month-old C57BL/6 WT mice and retinal flatmounts were made 1 and 2 months after the injection to observe AcGFP expression. The retinas at 1 month after the injection showed a strong AcGFP signal only around the AAV-injected area. The AcGFP region spread in the retinas at 2 months after injection (Fig. 4B) and increase in AcGFP-positive cells was observed in experimental retinas injected with AAV8-SpCas9, AAV8-Donor, and scAAV8-U6-gRNAx2, in which the number of AcGFP-positive cells was similar to that observed after in vivo electroporation. At 2 months in the retinas of the negative control group, sporadic AcGFP signals were detected around the injection area. Because the ITR (inverted terminal repeat) sequences of AAVs have transcriptional activity^39^, the AcGFP signals may indicate leakage of ITR-mediated transcription.

**Figure 4.**
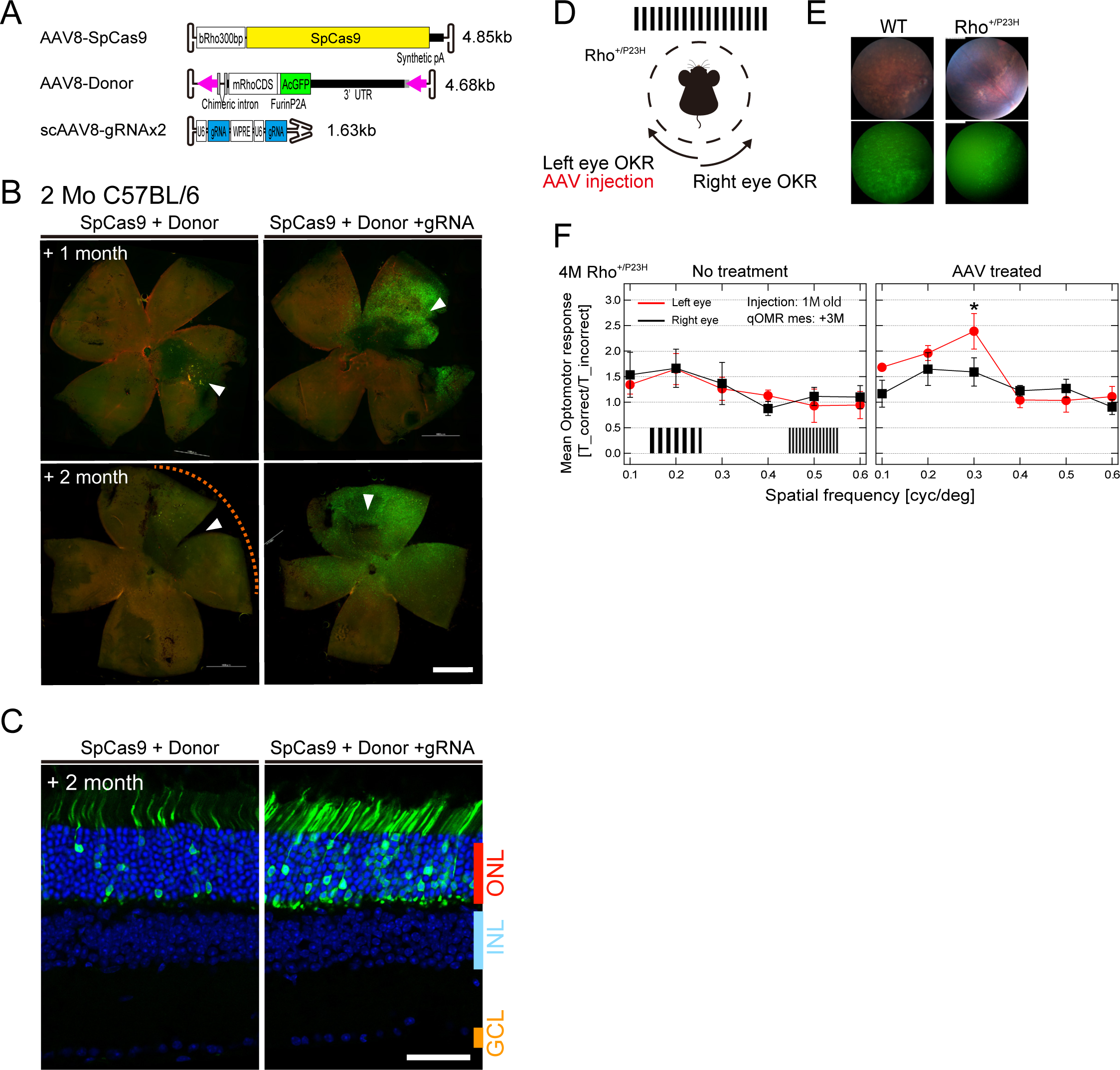
Effectiveness of HITI-treatment gene constructs packaged into AAV8 vectors. (A) Schematic illustration of the HITI-treatment AAV constructs. mRho-gRNA1 was used as the target. The nucleotide length of each AAV construct is shown on the right. (B) Flatmounts of subretinally injected retinas with (right) or without (left) mRho-gRNA1 expressing AAV (scAAV8-U6-gRNA×2). AAV cocktails were injected into 2-month-old C57BL/6J mice, and the retinas injected after 1 (upper) and 2 (lower) months are shown. Scale bar, 1 mm. (C) Immunohistochemistry sections of retinas at 2 months after injection immunostained with anti-GFP antibody. Scale bar, 50 μm. (D) Schematic illustration of measurement of the optomotor response (OMR). The OMRs from the left and right eyes were recorded using clockwise and counterclockwise moving patterns, respectively. The HITI-treatment AAV cocktail was injected into the left eye of 1 month Rho^+/P23H^ mice. (E) Fluorescent fundus image 2 months after injection. (F) OKR measurements in the left and right eye. The graph shows representative measurements from untreated (left) and AAV-treated (right) Rho^+/P23H^ mice. The horizontal and vertical axes represent the spatial frequency of the moving patterns and the OKR response, respectively. OKR was measured in the left (red) and right (black) eyes. Data represents means□±□SD (n = 3) with individual experimental points. *p□<□0.05

To evaluate the possible therapeutic effects of AAV in mutant mice, we examined the pattern perception of HITI-treated mice using the optomotor response (OMR), which is observed as the head movement of mice synchronous to moving patterns of various spatial frequencies. Because Rho^P23H/P23H^ homozygous mice (Fig. 3) showed complete degeneration of photoreceptor cells before AAV reconstitution, Rho^+/P23H^ heterozygous mice were used. AAV cocktails were injected into the left eye of 1-month-old mice (Fig. 4D), and 1 month after injection, mice with AcGFP-positive retinas were selected using fluorescent fundus imaging (Fig. 4G). The number of AcGFP-expressing cells was lower in these mice than in WT mice, possibly because of the progression of photoreceptor degeneration.

Two months after injection, the OMR was measured. Visual stimuli were displayed under a filter set for scotopic measurements, in which rod photoreceptors were evoked predominantly. In the WT mice, OMR-driven head movement was observed by moving patterns of spatial frequency between 0.1 c/deg and 0.3 c/deg, and Rho^+/P23H^ mice without AAV treatment responded to the moving patterns. In HITI-treated mice, the OMR response was significantly increased in the left eye, which suggested increased sensitivity of rod-derived perception because of HITI-mediated mRhoCDS expression instead of Rho^P23H^ expression.

### Rapid validation of highly efficient HITI-mediated gene constructs for the mouse *Prph2* locus

To examine whether our workflow for HITI-mediated gene construction can be applied to other genes, we targeted mouse Prph2, the second leading cause of AdRP^40,41^. In the grID database in silico finder, three gRNA target sequences with grID scores ranging from 800 to 900 were selected around the 100 bp region of the proximal sequence from the Prph2 5’-UTR (Fig. 5A). We then constructed plasmids for the HITI-mediated mPrph2CDS insertion targeting the three gRNA sequences and performed in vivo electroporation. In the donor plasmid, mRho cDNA was replaced with mPrph2 cDNA and the 3′-UTR was replaced with the Prph2 3′-UTR (Fig. 5A). In P21 mouse retinas, HITI constructs with mPrph2-gRNA1 and mPrph2-gRNA3 showed adequate AcGFP fluorescence signals (Fig. 5B, 5C). Most mCherry-positive cells in the ONL coexpressed AcGFP, which suggested that these HITI-treatment constructs were highly effective. However, the HITI construct with mPrph2-gRNA2 showed low EGFP expression even though the grID score was as high as 900.

**Figure 5.**
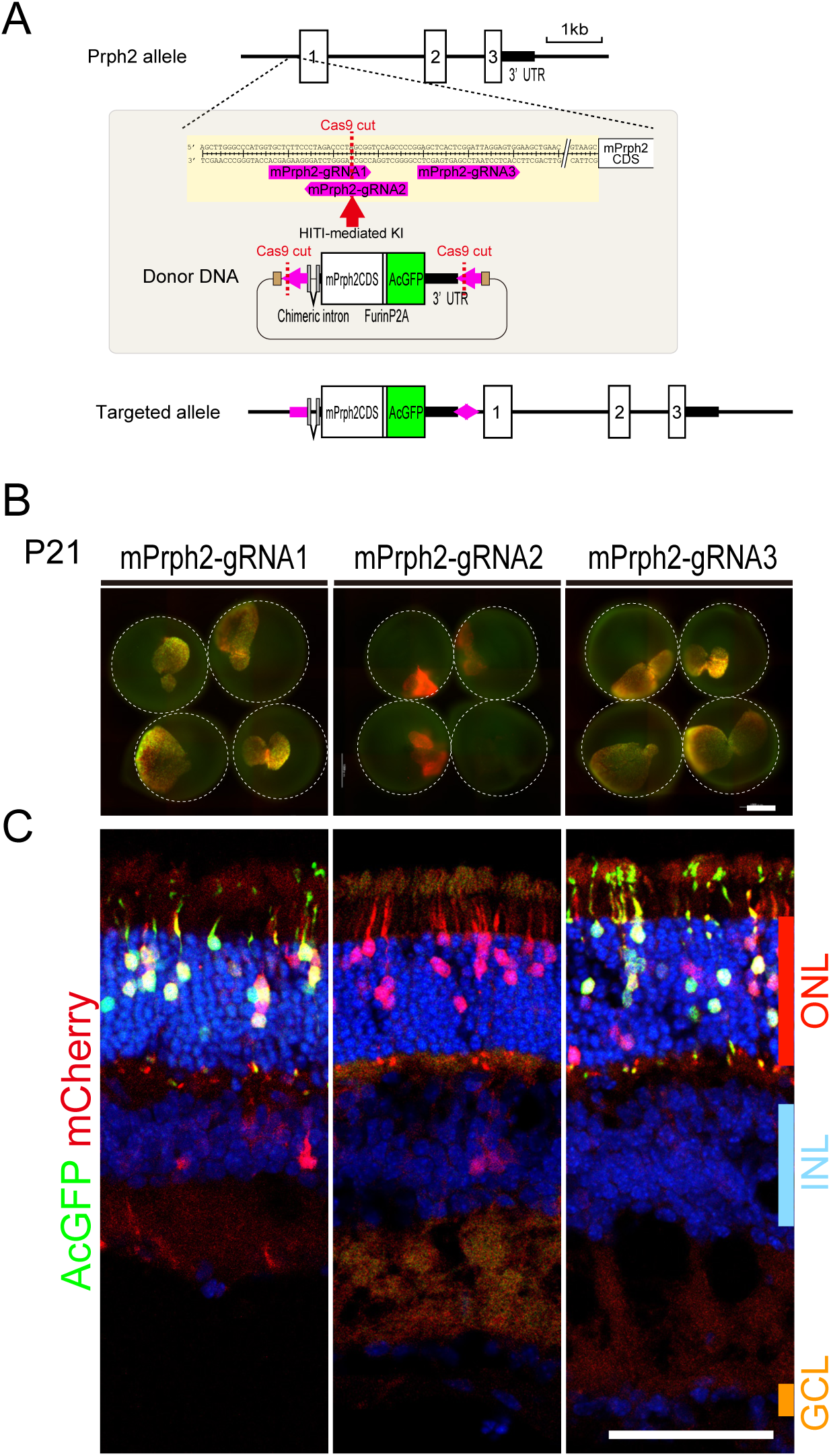
Optimization and validation of highly efficient HITI-treatment gene constructs targeting the mouse *Prph2* locus. (A) Schematic illustration of the Cas9-driven HITI-mediated gene insertion into the mouse *Prph2* locus. Each donor cassette containing one of the three gRNA sequences (pink) selected from the 5′-UTR sequence of the mouse *Prph2* locus at both ends was prepared. The targeted allele represents the gene structure when the HITI-mediated gene insertion occurs. (B) P21 mouse eyecups electroporated in vivo at P0 with three plasmid cocktails targeting the mPrph2-gRNA1 (left), mPrph2-gRNA2 (middle), and mPrph2-gRNA3 (right) sequences of the mouse *Prph2* locus. The dashed circles represent each eyecup. Scale bar, 1 mm. (C) Immunohistochemical analysis of HITI-treated P21 mouse retinas targeting three mPrph2 gRNAs. Electroporated cells were mCherry positive, and most cells in the ONL expressed AcGFP in retinal sections targeting gRNA and gRNA3 sequences. Scale bar, 50 μm.

## Discussion

The advantage of the HITI-based genome-editing technique is the ability to insert exogenous DNAs in the desired orientation at the target site in nondividing cells^19,20,34^. In in vivo gene therapy, most target cells are differentiated, that is, the cells are in a nondividing state. The insertion of exogenous genes is mediated by NHEJ and/or homologous-directed repair (HDR), which are cellular genome repair machineries triggered by DSBs. Because HDR occurs less frequently in nondividing cells, the HDR-mediated in vivo exogenous gene insertion efficiency in mouse retinas is about 10%^42^. Given that HITI gene insertion is mediated by NHEJ, the gene insertion efficiency of differentiated mouse rod photoreceptors can be as high as 80%–90%. The efficiency observed in this study was also higher than that in previous studies targeting the mouse Rho gene^21,22^ possibly because of the use of a *Rho* promoter, which is highly active in rod photoreceptor cells.

With *Rho* promoters, HITI-mediated gene insertion is facilitated by efficient cleavage of the target genome and donor vector. Evaluation of the manner of insertion in single-rod photoreceptor cells indicated that 22% were biallelic and 78% were monoallelic. The HITI-mediated insertion was similar to that observed in the HITI experiment using Rho^+/AcGFP^ KI mice. Because the ratio of monoallelic insertions to the AcGFP KI allele was included in the AcGFP-negative fraction, the percentage of gene insertions was higher for the WT allele than for *AcGFP* allele, possibly because of the difference in genome structure between the WT and *AcGFP* alleles. It is possible that the donor cassette was difficult to insert because *AcGFP* had already been inserted.

Most mutations in *RHO* are dominant inheritance mechanisms that cause dominant inhibition or haploinsufficiency^23,24^; thus, the syndrome emerges in heterozygotes. In this study, the percentages of monoallelic and biallelic HITI-mediated mRhoCDS insertions were 78% and 22%, respectively. Based on these results, about 60% of HITI-treated rod photoreceptor cells were expected to be repaired when HITI was performed in mutant heterozygote individuals. That is, assuming that monoallelic insertion occurs equally in the mutant and normal *RHO* alleles, normal *RHO* was expressed for 22% of biallelic insertions and 39% of monoallelic insertions.

It is essential that the expression level of the inserted gene is restored to the same level as that of the WT to achieve high therapeutic efficacy in treating rhodopsin mutations. Therefore, we optimized the construction of the donor cassette. When generating KI or transgenic mice expressing exogenous cDNA, inserting an intron before the cDNA increases the transcription level. Because 95% of the proteins expressed in rod outer segments are rhodopsins^27^, a high expression level must be maintained. It was previously reported that the transcription and expression levels of KI genes increase upon insertion of chimeric intron into the 5′-UTR of the mouse *Rho* allele^31,32^. The 3′-UTR region of the mouse *Rho* gene should be considered because this gene produces multiple mRNA transcripts that affect RNA transport and stability^33^. We used the chimeric intron and 3′-UTR in the donor cassette, and HITI-treated rod photoreceptors in Rho^P23H/P23H^ mutant mice were maintained with elongated outer segments. This result indicated that normal rhodopsin proteins expressed in optimized donor cassettes were sufficient to suppress photoreceptor degeneration.

HITI-treated gene constructs can be used as AAV vectors. The insertion efficiency was similar to that for in vivo electroporation. In addition, AAV-treated Rho^+/P23H^ mutant mice showed good recovery in OMR measurements. Rho^+/P23H^ heterozygous mice did not exhibit progressive degeneration, but rod photoreceptor sensitivity was lower than in WT mice^38^. This recovery could be attributed to the increased sensitivity of HITI-treated photoreceptor cells expressing normal rhodopsins. In HITI-treated rod photoreceptors, the remaining mutant genome is considered to be activated. Considering that AcGFP expression was stopped by the HITI-mediated insertion of the mCherry-containing donor cassette into the Rho^+/AcGFP^ mouse retinas, the remaining mutant gene was not expressed.

In this study, we established a workflow for the rapid validation of highly efficient HITI-treatment gene constructs, which is expected to contribute to the efficient development of gene therapy products in the ophthalmology field, especially for photoreceptor cells, and to shorten the development time. When SpCas9 is used as a genome-editing tool, three gRNA sequences selected by the in silico gRNA finder would be sufficient for determining a good gRNA sequence for in vivo electroporation, although there is no significant correlation between in silico scores and the cleavage efficiency tested in vitro and in vivo. Testing this initial validation using AAV would be time and cost intensive. Plasmids with appropriate backbones are key to successful HITI-mediated insertion via in vivo electroporation. In pLeaklessIII plasmids, triple poly(A) sequences are inserted into donor cassettes to prevent anomalous transcription from the plasmid backbone^34^. Switching the donor plasmid backbone to pAAV resulted in no HITI-mediated donor cassette insertion because there would be transcriptional activity from both the plasmid backbone and AAV ITR sequence^39^. It is likely that the HITI-treated AAV vectors functioned efficiently in our study because the pAAV plasmid backbone had been removed. If such undesired transcription inhibits the HITI machinery, insertion of poly(A) between the ITR and donor cassettes may improve HITI efficiency.

We validated the HITI-treatment gene constructs for the mouse *Prph2* locus from two of three gRNA target sequences. It took about 2 months for vector construction and in vivo validation, which shows that the workflow is convenient for application to other AdRP target genes. The workflow developed in this study can also be applied to human systems. For example, humanized mice carrying partially human genomic sequences can be used for in vivo validation, and electroporation-based gene transfer can be tested with human stem cell-derived organoids. This time- and cost-effective workflow may help to accelerate the development of HITI-based genome-editing reagents.

## Experimental Procedures

### Plasmids

The details of the DNA constructs used in this study are listed in Table S1.

### Mice

All mouse experiments were conducted with approval from the RIKEN Center for Developmental Biology Ethics Committee (No. AH18-05-23). Timed pregnant CD1 (Charles River Laboratories, Japan) and C57BL/6J (CLEA, Japan) mice were maintained and provided by the Laboratory for Animal Resources and Genetic Engineering (RIKEN Biosystems Dynamics Research (BDR)). Rho^P23H^ mutant mice were purchased from Jackson Laboratory. Rho^AcGFP^ KI mice were generated by the Laboratory for Animal Resources and Genetic Engineering, RIKEN BDR, according to a previously reported procedure^43^. Briefly, the donor vector had 5′- (1000 bp) and 3′- (1000 bp) homology arms for intron 2 of the mouse *Rho* locus and an expression cassette containing an adenovirus splicing acceptor, stop codons, an internal ribosome entry site, AcGFP, and a rabbit beta-globin poly(A) sequence. crRNAs for intron 2 (mRhoIntron2-crRNA; 5′-UAG AGA GCA UUG CCG UUA CUG UUU UAG AGC UAU GCU GUU UUG-3′) and tracrRNA (5′-AAA CAG CAU AGC AAG UUA AAA UAA GGC UAG UCC GUU AUC AAC UUG AAA AAG UGG CAC CGA GUC GGU GCU-3′) were purchased (FASMAC, Japan). Zygotes generated by in vitro fertilization were microinjected with 100 ng/μL Cas9 protein (Thermo Fisher Scientific), 50 ng/μL mRhoIntron2-crRNA, 100 ng/μL tracrRNA, and 10 ng/μL donor vector.

### Single-strand annealing assay

HEK293T cells were maintained in DMEM (high glucose, Fujifilm Wako) supplemented with 10% FBS. Plasmids of 100 ng of pCAG-SpCas9, 100 ng of pBAsi-U6-mRho-gRNA, 300 ng of pCAG-EGxxFP, and 100 ng of pCAG-mCherry were transfected into 80% confluent HEK293T cells in 24-well plates using FuGene6 (Roche) according to the manufacturer’s protocol. After 48 h of incubation, EGFP and mCherry expression was observed using a fluorescence microscope, and cleavage efficiency was evaluated by EGFP:mCherry ratios calculated from Pixel intensity values of GFP-positive and mCherry-positive areas measured by ImageJ.

### In vivo electroporation and eyecup preparation

In vivo electroporation was performed at P0, as previously described with minor modifications^44^. In brief, 0.4 μL of plasmid DNA solutions of 0.16 μg/μL pRho2k/300bp-SpCas9, 0.16 μg/μL pLeaklessIII/pAAVV donor cassettes, 0.06 μg/μL pBAsi-U6-mRho/mPrph2-gRNAs, and 0.02 μg/μL pCAG-mCherry were injected into the subretinal space of P0 mouse pups with a 33G blunt-ended microsyringe (ITO CORPORATION) and transferred into the retina by an NEPA21 Super Electroporator (Nepagene) with tweezer-type electrodes (Nepagene, CUY650P10). The parameters were as follows: voltage, 120 V; pulse length, 30 ms; pulse interval, 470 ms; number of pulses, 3; decay rate, 10%; polarity + as poring pulse; and voltage, 20 V; pulse length, 50 ms; pulse interval, 50 ms; number of pulses, 3; decay rate, 40%; and polarity + as transfer pulse. Electroporated retinas were harvested at P14, P21, P50, and/or P56 for IHC analysis.

### AAV construction and subretinal injection

pAAV or pscAAV plasmids carrying the SpCas9, donor cassettes, or gRNAs listed in Supplemental Table S1 were constructed. All AAV constructs were packaged into AAV8 serotypes by the Gene Transfer, Targeting, and Therapeutics Viral Vector Core of the Salk Institute or VectorBuilder. Subretinal injection of AAVs was performed as previously described with minor modifications^45^. In brief, 1- or 2-month-old C57BL/6J and Rho^+/P23H^ mice were anesthetized with ketamine (7.7 mg/100 g; Daiichi Sankyo) and xylazine (0.92 mg/100 g; Bayer), and 1 μL of 2.0 × 10^12^ vg/mL AAV solution of 0.8 × 10^12^ vg/mL AAV8-300bp-SpCas9, 0.8 × 10^12^ vg/mL AAV8-donor cassettes, or 0.4 × 10^12^ vg/mL AAV8-U6-gRNAs was injected into the subretinal space. Retinas were harvested 1 and 2 months after injection.

### Section immunohistochemistry and retinal flatmounts

Retinal eyecups, in which the sclera and cornea were dissected, were fixed with 4% paraformaldehyde in PBS (Nacalai Tesque, Inc.) for 1 h at room temperature. After washing with PBS, mCherry- and/or AcGFP-positive eyecups were used for section IHC and retinal flatmounts. Immunostaining and imaging of the retinal sections^44^ and flatmounts^46^ were performed as previously described. Primary antibodies used in this study were directed against GFP (rat, 1:1000, Nacalai Tesque, Inc., 04404-84), mCherry (rabbit, 1:1000, MBL, PM-005), and rhodopsin (mouse, 1:1000, Merck, MAB5316). Fluorescence images were acquired using a confocal microscope (LSM 700, Zeiss) and a BZ-9000 fluorescence microscope (Keyence). To count AcGFP- and mCherry-positive cells, four to six images of a 320 μm × 320 μm field were obtained using confocal microscopy.

### Single cell sorting and genotyping

The P56 CD1 retinas were electroporated in vivo with plasmids containing SpCas9. Donor vectors carrying NLS-AcGFP, U6-gRNA, and NLS-mCherry were dissected to isolate fluorescent retinal regions and were dissociated into single cells with papain digestion as previously described^47^. The dissociated cells were centrifuged, resuspended in PBS containing 0.5% bovine serum albumin, and filtered through a 35 μm mesh (352235, Coning). The GFP- and mCherry-double-positive cells were directly sorted into 2 μL of cell lysis buffer in a 96-well PCR plate using an SH800 cell sorter and SH800 software (version 2.1.5, Sony), according to the manufacturer’s recommendations.

Using the lysed cells as the templates, the first PCR was performed with primer pairs 5′-GCAGCAGTGGGATTAGCGTTAGTATG-3′ and 5′-AAGGGCACATAAAAATTGGGGCCCTC-3′ to amplify both the WT (350 bp) and KI (about 600 bp) fragments. Using 1:200 diluted PCR products, a second PCR (nested PCR) was performed with primer pairs to amplify the WT allele (5′-TATCTCGCGGATGCTGAATCAGCCTC-3′ and 5′-AAGGGCACATAAAAATTGGGGCCCTC-3′) and the KI allele (5′-TATCTCGCGGATGCTGAATCAGCCTC-3′ and 5′-TTCTCTGTCTCGACAAGCCCAGTTTC-3′). For direct sequencing, PCR products that were reamplified using nested PCR were collected and used. All PCR analyses were performed using PrimeStarGXL (TaKaRa) as follows: 94°C for 10 min and 30 cycles of 98°C for 5 s, 55°C for 15 s, and 68°C for 30 s.

### Quantitative optomotor response

A quantitative optomotor system (Phenosys GmbH, Berlin, Germany) was used to determine the visual acuity according to the manufacturer’s instructions. Briefly, under an ND2 filter set for scotopic measurements, sinusoidal gratings ranging from 0.1 to 0.6 cycles per degree and at maximal contrast^48^ were displayed in randomized order for 1 min. Data were averaged from three sessions per time point for each mouse using integrated software for the detection of head-tracking movements and projection of a virtual cylinder of revolving gratings of different spatial frequencies. Each mouse was tested for no more than 30 min.

## Acknowledgments

We received generous support from all members of the Laboratory of Retinal Regeneration, RIKEN Center for Biosystems Dynamics Research. We thank the members of the Laboratory for Animal Resources and Genetic Engineering, RIKEN Center for Biosystems Dynamics Research, for providing and maintaining the mice and H. Fujiwara for supporting the FACS analyses. This work was supported in part by grants from the Japan Agency for Medical Research and Development (grant number 17bm0204002h0005 to M.T.), JSPS KAKENHI (grant numbers 24687010 and 17K11471 to A.O.), and the Charitable Trust Fund for Ophthalmic Research in the Commemoration of Santen Pharmaceutical’s Founder (to A.O.).

## Ethics

All animal experiments were conducted with approval from the RIKEN Center for Developmental Biology Ethics Committee (No. AH18-05-23).

---

**Supplemental Figure S1.** P21 mouse eyecups were electroporated in vivo at P0 using three plasmid cocktails targeting different gRNA1 (left), gRNA2 (middle), and gRNA3 (right) sequences. The dashed circles represent each eyecup. Scale bar, 1 mm.

**Supplemental Figure S2. Single-cell genotyping by nested PCR**

FACS was used to sort 48 GFP- and mCherry-double-positive cells (Fig. 2B). Electrophoresis gel images of nested PCR of the WT and KI alleles are shown. Of the 48 samples, nine were biallelic and showed only KI PCR bands, and 28 were monoallelic and showed both WT and KI PCR bands.

**Supplemental Figure S3. Generation of Rho^AcGFP^ knock-in mice**

A schematic of this process is shown. The donor vector carrying the AcGFP expression cassette containing an adenovirus splicing acceptor, stop codons, an internal ribosome entry site, AcGFP, and a rabbit beta-globin polyadenylation (pA) sequence was inserted into intron 2, and its expression was triggered by Cas9-driven cleavage DS (B).

**Supplemental Figure S4. Ineffectiveness of the AAV vector integrated the donor and U6-gRNA cassettes**

(A) Schematic illustration of the HITI-treatment AAV constructs. The nucleotide length of each AAV construct is shown on the right.

(B) Eyecups 1 month after subretinal injection with AAV cocktails in 2-month-old C57BL/6J mice. AAV cocktails were injected with (right) and without (left) SpCas9 expressing AAV (AAV8-SpCas9). Scale bar, 1 mm.

## References

1. Grover, S. et al. Visual acuity impairment in patients with retinitis pigmentosa at age 45 years or older. Ophthalmology 106, 1780–1785 (1999).

2. O’Neal, T. B. & Luther, E. E. Retinitis Pigmentosa. StatPearls (2023).

3. Hamel, C. Retinitis pigmentosa. Orphanet J Rare Dis 1, 40 (2006).

4. Verbakel, S. K. et al. Non-syndromic retinitis pigmentosa. Prog Retin Eye Res 66, 157–186 (2018).

5. Pontikos, N. et al. Genetic basis of inherited retinal disease in a molecularly characterized cohort of more than 3000 families from the United Kingdom. Ophthalmology 127, 1384–1394 (2020).

6. Koyanagi, Y. et al. Genetic characteristics of retinitis pigmentosa in 1204 Japanese patients. J Med Genet 56, 662–670 (2019).

7. Schneider, N. et al. Inherited retinal diseases: Linking genes, disease-causing variants, and relevant therapeutic modalities. Prog Retin Eye Res 89, 101029 (2022).

8. Arabi, F., Mansouri, V. & Ahmadbeigi, N. Gene therapy clinical trials, where do we go? An overview. Biomed Pharmaco 153, 113324 (2022).

9. Chiu, W. et al. An update on gene therapy for inherited retinal dystrophy: Experience in Leber congenital amaurosis clinical trials. Int J Mol Sci, 22, 4534 (2021).

10. Hu, M. L. et al. Gene therapy for inherited retinal diseases: Progress and possibilities. Clin Exp Optom 104, 444–454 (2021).

11. Cross, N., van Steen, C., Zegaoui, Y., Satherley, A. & Angelillo, L. Current and future treatment of retinitis pigmentosa. Clin Ophthalmol 16, 2909 (2022).

12. Toualbi, L., Toms, M. & Moosajee, M. The landscape of non-viral gene augmentation strategies for inherited retinal diseases. Int J Mol Sci, 22, 2318 (2021).

13. Daich Varela, M., Georgiadis, A. & Michaelides, M. Genetic treatment for autosomal dominant inherited retinal dystrophies: Approaches, challenges and targeted genotypes. Br J Ophthalmol 107, 1223–1230 (2023).

14. van der Oost, J. & Patinios, C. The genome editing revolution. Trends Biotechnol 41, 396–409 (2023).

15. Wang, J. Y. & Doudna, J. A. CRISPR technology: A decade of genome editing is only the beginning. Science 379, 6629 (2023).

16. Saber Sichani, A., Ranjbar, M., Baneshi, M., Torabi Zadeh, F. & Fallahi, J. A review on advanced CRISPR-based genome-editing tools: base editing and prime editing. Mol Biotechnol 65, 849–860 (2023).

17. Botto, C., Dalkara, D. & El-Amraoui, A. Progress in gene editing tools and their potential for correcting mutations underlying hearing and vision loss. Front Genome Ed 3, 737632 (2021).

18. Diakatou, M., Manes, G., Bocquet, B., Meunier, I. & Kalatzis, V. Genome editing as a treatment for the most prevalent causative genes of autosomal dominant retinitis pigmentosa. Int J Mol Sci 20, (2019).

19. Suzuki, K. & Izpisua Belmonte, J. C. In vivo genome editing via the HITI method as a tool for gene therapy. J Human Genet 63, 157–164 (2017).

20. Suzuki, K. et al. In vivo genome editing via CRISPR/Cas9 mediated homology-independent targeted integration. Nature 540, 144–149 (2016).

21. Tornabene, P. et al. Therapeutic homology-independent targeted integration in retina and liver. Nat Commun 13, 1–14 (2022).

22. Hoang, D. A., Liao, B., Zheng, Z. & Xiong, W. Mutation-independent gene knock-in therapy targeting 5′UTR for autosomal dominant retinitis pigmentosa. Sig Transduct Target Ther 8, 1–3 (2023).

23. Athanasiou, D. et al. The molecular and cellular basis of rhodopsin retinitis pigmentosa reveals potential strategies for therapy. Prog Retin Eye Res 62, 1 (2018).

24. Zhen, F. et al. Rhodopsin-associated retinal dystrophy: Disease mechanisms and therapeutic strategies. Front Neurosci 17, 1132179 (2023).

25. Tebbe, L., Kakakhel, M., Makia, M. S., Al-Ubaidi, M. R. & Naash, M. I. The interplay between peripherin 2 complex formation and degenerative retinal diseases. Cells 2020, 9, 784 (2020).

26. Jaskula-Ranga, V. & Zack, D. J. grID: A CRISPR-Cas9 guide RNA database and resource for genome-editing. bioRxiv 097352 (2016) doi:10.1101/097352.

27. McDowell, J. H. Preparing rod outer segment membranes, regenerating rhodopsin, and determining rhodopsin concentration. Methods in Neurosciences 15, 123–130 (1993).

28. Brinster, R. L., Allen, J. M., Behringer, R. R., Gelinas, R. E. & Palmiter, R. D. Introns increase transcriptional efficiency in transgenic mice. Proc Natl Acad Sci U S A 85, 836–840 (1988).

29. Buchman, A. R. & Berg, P. Comparison of intron-dependent and intron-independent gene expression. Mol Cell Biol 8, 4395–4405 (1988).

30. Choi, T., Huang, M., Gorman, C. & Jaenisch, R. A generic intron increases gene expression in transgenic mice. Mol Cell Biol 11, 3070–3074 (1991).

31. Sakurai, K. et al. Physiological properties of rod photoreceptor cells in green-sensitive cone pigment knock-in mice. J Gen Physiol 130, 21–40 (2007).

32. Kodama, T. et al. Expression and localization of an exogenous G protein-coupled receptor fused with the rhodopsin C-terminal sequence in the retinal rod cells of knockin mice. Exp Eye Res 80, 859–869 (2005).

33. Al-Ubaidi, M. R. et al. Mouse opsin. Gene structure and molecular basis of multiple transcripts. J Biol Chem 265, 20563–20569 (1990).

34. Tsunekawa, Y. et al. Developing a *de novo* targeted knock-in method based on *in utero* electroporation into the mammalian brain. Development 143, 3216–3222 (2016).

35. Tsunekawa, Y., Terhune, R. K. & Matsuzaki, F. Protocol for de novo gene targeting via in utero electroporation. Methods Mol Biol 2312, 309–320 (2021).

36. Matsuda, T. & Cepko, C. L. Controlled expression of transgenes introduced by *in vivo* electroporation. Proc Natl Acad Sci U S A 104, 1027–1032 (2007).

37. Sakami, S. et al. Probing mechanisms of photoreceptor degeneration in a new mouse model of the common form of autosomal dominant retinitis pigmentosa due to P23H opsin mutations. J Biol Chem 286, 10551–10567 (2011).

38. Sakami, S., Kolesnikov, A. V., Kefalov, V. J. & Palczewski, K. P23H opsin knock-in mice reveal a novel step in retinal rod disc morphogenesis. Hum Mol Genet 23, 1723–1741 (2014).

39. Robichaux, M. A. et al. Subcellular localization of mutant P23H rhodopsin in an RFP fusion knock-in mouse model of retinitis pigmentosa. Dis Model Mech 15, dmm049336 (2022).

40. Haberman, R. P., McCown, T. J. & Samulski, R. J. Novel transcriptional regulatory signals in the adeno-associated virus terminal repeat A/D junction element. J Virol 74, 8732 (2000).

41. Peeters, M. H. C. A., et al. *PRPH2* mutation update: In silico assessment of 245 reported and 7 novel variants in patients with retinal disease. Hum Mutat 42, 1521–1547 (2021).

42. Oishi, A. et al. Genetic and phenotypic landscape of *PRPH2*-associated retinal dystrophy in Japan. Genes 12, 1817 (2021).

43. Matsuda, T. & Oinuma, I. Optimized CRISPR/Cas9-mediated in vivo genome engineering applicable to monitoring dynamics of endogenous proteins in the mouse neural tissues. Sci Rep 9, 11309 (2019).

44. Abe, T., Inoue, K.-I., Furuta, Y. & Kiyonari, H. Pronuclear microinjection during S-phase increases the efficiency of CRISPR-Cas9-assisted knockin of large DNA donors in mouse zygotes ll. (2020) doi:10.1016/j.celrep.2020.107653.

45. Onishi, A. et al. Pias3-dependent SUMOylation directs rod photoreceptor development. Neuron 61, 234–246 (2009).

46. Assawachananont, J. et al. Transplantation of embryonic and induced pluripotent stem cell-derived 3D retinal sheets into retinal degenerative mice. Stem Cell Rep 2, 662–674 (2014).

47. Onishi, A., Peng, G. H., Chen, S. & Blackshaw, S. Pias3-dependent SUMOylation controls mammalian cone photoreceptor differentiation. Nat Neurosci 13, 1059–1065 (2010).

48. Maekawa, Y. et al. Optimized culture system to induce neurite outgrowth from retinal ganglion cells in three-dimensional retinal aggregates differentiated from mouse and human embryonic stem cells. Curr Eye Res 41, 558–568 (2016).

49. Kretschmer, F., Tariq, M., Chatila, W., Wu, B. & Badea, T. C. Comparison of optomotor and optokinetic reflexes in mice. J Neurophysiol 118, 300–316 (2017).

