## Supplemental Tables and Figures for "Efficient workflow for validating homology-independent targeted integration-mediated gene insertion in rod photoreceptor cells to treat dominant-negative mutations causing retinitis pigmentosa"

### Supplemental Table S1 DNA construction

#### DNAs

| Description in the text | cloned DNA/genes | cloning method |
|---|---|---|
| EGFP fragment for EGxxFP | EGFP (1-600bp) with stop codon, EGFP (120-720bp) | PCR with PrimerStarGXL (TaKaRa)*1 |
| mRho gRNA target | 5'-CTGAGCTCGCCAAGCAGCCTTGGTCTCTGTCTACGAAGAGCCCGTGGGGCAGCCTCGAGAGCC-3' | Oligonucleotide synthesis |
| mRho-gRNA1 | 5'-TCTGTCTACGAAGAGCCCGTGGG-3' (grID: mm106628048, grID score = high[800])*2 | Oligonucleotide synthesis |
| mRho-gRNA2 | 5'-GGCTCTCGAGGCTGCCCCACGGG-3' (grID: mm106628053, grID score = high[800])*2 | Oligonucleotide synthesis |
| mRho-gRNA3 | 5'-CTGAGCTCGCCAAGCAGCCTTGG-3' (grID: mm106628045, grID score = high[600])*2 | Oligonucleotide synthesis |
| gRNA scaffold | 5'-ATAGCAAGTTAAAATAAGGCTAGTCCGTTATCAACTTGAAAAAGTGGCACCGAGTCGGTGC-3' | Oligonucleotide synthesis |
| NLS | nuclear localization signal from c-Myc (PAAKRVKLD) | Oligonucleotide synthesis |
| bRho2k | bovine rhodopsin 2k promoter (chr22:56,231,474-56,233,726 in ARS-UCD1.2/bosTau9) | PCR with PrimerStarGXL (TaKaRa)*1 |
| bRho300bp | bovine rhodopsin 300bp promoter (chr22:56,231,474-56,231,769 in ARS-UCD1.2/bosTau9) | PCR with PrimerStarGXL (TaKaRa)*1 |
| SpCas9 | CDS of Streptococcus pyogenes Cas9 | PCR with PrimerStarGXL (TaKaRa)*1 |
| synthetic pA | Synthetic polyadelylation signal (5'-AATAAAAGATCTTTATTTTCATTAGATCTGTGTGTTGGTTTTTTGTGTG-3') | Oligonucleotide synthesis |
| Chimeric Intron | Chimeric intron of human beta globin and IgG | Oligonucleotide synthesis |
| mRhoCDS | mouse Rhodopsin CDS (NM_145383.2) | PCR with PrimerStarGXL (TaKaRa)*1 |
| FurinP2A | Furin and P2A self digestive peptides with GSG linker (RRKR-GSG-ATNFSLLKQAGDVEENPGP) | Oligonucleotide synthesis |
| mRho3'UTR | 3'UTR from mouse rhodopsin locus (chr6:115,936,720-115,938,976 in GRCm38/mm10) | PCR with PrimerStarGXL (TaKaRa)*1 |
| mPrph2-gRNA1 | 5'-TGCTCTTCCCTAGACCCTAGCGG-3' (grID: mm056084424, grID score = high[800])*2 | Oligonucleotide synthesis |
| mPrph2-gRNA2 | 5'-GGGCTGGACCGCTAGGGTCTAGG-3' (grID: mm056084433, grID score = high[900])*2 | Oligonucleotide synthesis |
| mPrph2-gRNA3 | 5'-GAGCTCACTCGGATTAGGAGTGG-3' (grID: mm056084434, grID score = high[800])*2 | Oligonucleotide synthesis |
| mPrph2CDS | mouse Peripherin2 CDS (NM_008938.2) | PCR with PrimerStarGXL (TaKaRa)*1 |
| mPrph3'UTR | 3'UTR from mouse Prph2 locus (chr17:46,923,548-46,925,033 in GRCm38/mm10) | PCR with PrimerStarGXL (TaKaRa)*1 |
| mRho 5' franking arm | chr6:115,933,764-115,934,808 in GRCm38/mm10 | PCR with PrimerStarGXL (TaKaRa)*1 |
| mRho 3' franking arm | chr6:115,934,830-115,935,838 in GRCm38/mm10 | PCR with PrimerStarGXL (TaKaRa)*1 |

#### Plasmids

| Description in the text | closed DNA casettes (5' to 3' direction) | cloning method | Plasmid backbone |
|---|---|---|---|
| pCAG-SpCas9 | SpCas9 | InFusion HD Cloning Plus system (TaKaRa)*1 | pCAG (Addgene 11159) |
| pCAG-EGxxFP | [EGFP (1-600bp) with stop codon]-[mRho gRNA target]-[EGFP (120-720bp)] | InFusion HD Cloning Plus system (TaKaRa)*1 | pCAG (Addgene 11159) |
| pCAG-mCherry | mCherry | InFusion HD Cloning Plus system (TaKaRa)*1 | pCAG (Addgene 11159) |
| pCAG-NLS-mCherry | NLS, mCherry | InFusion HD Cloning Plus system (TaKaRa)*1 | pCAG (Addgene 11159) |
| pbRho2k-SpCas9 | bRho2k, SpCas9, synthetic pA | InFusion HD Cloning Plus system (TaKaRa)*1 | pCAG (Addgene 11159) |
| pbRho300-SpCas9 | bRho300bp, SpCas9, synthetic pA | InFusion HD Cloning Plus system (TaKaRa)*1 | pCAG (Addgene 11159) |
| pBAsi-U6-mRho-gRNA | mRho-gRNA(1/2/3)*3, gRNA scaffold | InFusion HD Cloning Plus system (TaKaRa)*1 | pBASi (TaKaRa) |
| pBAsi-U6-mPrph2-gRNA | mPrph2-gRNA(1/2/3)*3, gRNA scaffold | InFusion HD Cloning Plus system (TaKaRa)*1 | pBASi (TaKaRa) |
| (Illustrated in Fig.1C) | [mRho-gRNA(1/2/3)]*5-[ChimericIntron, mRhoCDS, FurinP2A, AcGFP, mRho3'UTR]-[mRho-gRNA(1/2/3)]*5 | InFusion HD Cloning Plus system (TaKaRa)*1 | pLeaklessIII*4 |
| (Illustrated in Fig.2A) | [mRho-gRNA1]*5-[ChimericIntron, mRhoCDS, FurinP2A, NLS, AcGFP, mRho3'UTR]-[mRho-gRNA1]*5 | InFusion HD Cloning Plus system (TaKaRa)*1 | pLeaklessIII*4 |
| (Illustrated in Fig.2E) | [mRho-gRNA1]*5-[ChimericIntron, mRhoCDS, FurinP2A, mCherry, mRho3'UTR]-[mRho-gRNA1]*5 | InFusion HD Cloning Plus system (TaKaRa)*1 | pLeaklessIII*4 |
| (Illustrated in Fig.5A) | [mPrph2-gRNA(1/2/3)]*5-[ChimericIntron, mPeph2CDS, FurinP2A, AcGFP, mPrph3'UTR]-[mPrph2-gRNA(1/2/3)]*5 | InFusion HD Cloning Plus system (TaKaRa)*1 | pLeaklessIII*4 |
| AAV8-SpCas9 | bRho300bp, SpCas9, synthetic pA | InFusion HD Cloning Plus system (TaKaRa)*1 | pAAV*6 |
| AAV8-Donor+gRNA | [U6-mRho-gRNA1, gRNA scaffold]-[mRho-gRNA1]*5-[ChimericIntron, mRhoCDS, FurinP2A, AcGFP, mRho3'UTR]-[mRho-gRNA1]*5 | InFusion HD Cloning Plus system (TaKaRa)*1 | pAAV*6 |
| AAV8-Donor | [mRho-gRNA1]*5-[ChimericIntron, mRhoCDS, FurinP2A, AcGFP, mRho3'UTR]-[mRho-gRNA1]*5 | InFusion HD Cloning Plus system (TaKaRa)*1 | pAAV*6 |
| scAAV8-U6-gRNA×2 | [U6-mRho-gRNA1, gRNA scaffold]-[WPRE]-[U6-mRho-gRNA1, gRNA scaffold] | InFusion HD Cloning Plus system (TaKaRa)*1 | pscAAV |
| (described in Fig.S4) | [mRho 5' franking arm]-[loxP]-[stop codon]-[IRES]-[AcGFP]-[b globin polyA]-[loxP]-[mRho 3' franking arm] *7 | InFusion HD Cloning Plus system (TaKaRa)*1 | pBluescriptII SK- |

*1 The reagents were used according to the manufacturer’s instructions.
*2 grID database (bioRxiv 097352; doi: https://doi.org/10.1101/097352)
*3 PAM sequences were removed.
*4 Development. 2016 Sep 1; 143(17): 3216–3222.
*5 these DNAs were inserted by reverse direction.
*6 The 5' AAV ITR sequence bears a 11-bp deletion (5'-AAAGCCCGGGC-3').
*7 This plasmid is used for generation of Rho<AcGFP> knock-in mice addressed in Supplemental Fig.2.

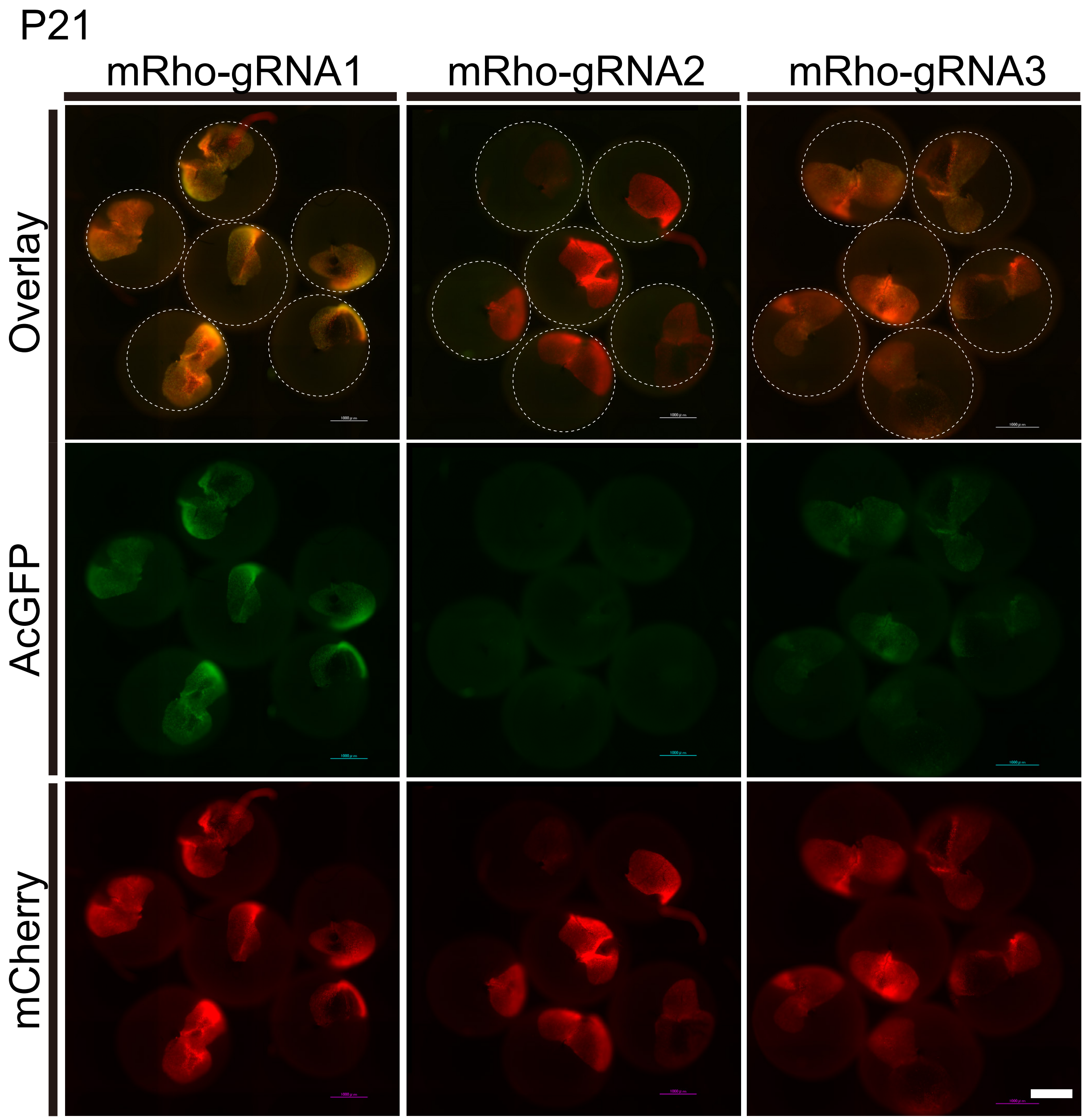


Supplemental Figure S1. Onishi et al.

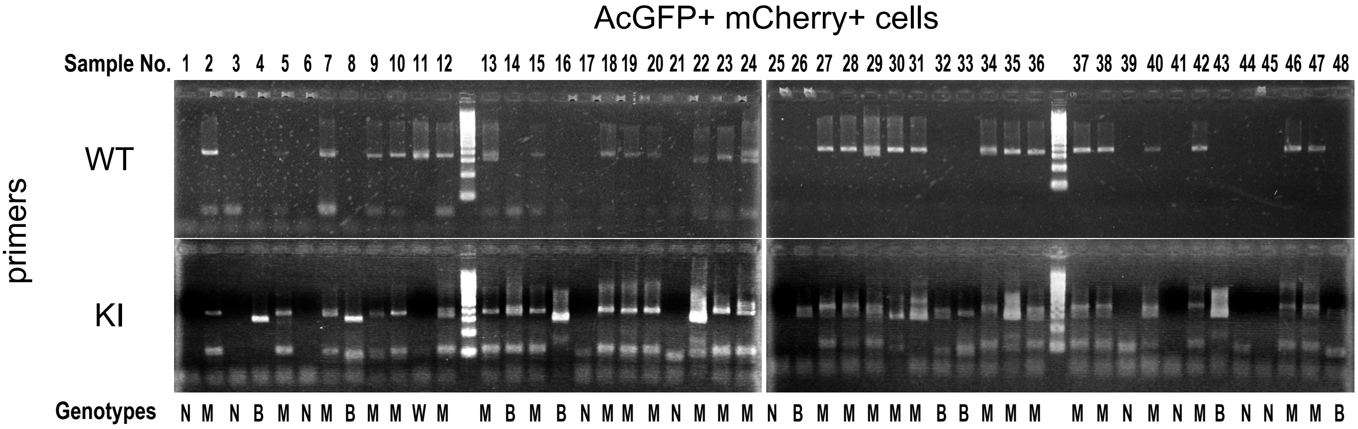


| | |
|---|---|
| B; biallelic | 9 |
| M; monoallelic | 28 |
| W; wildtype | 1 |
| N; not collected | 10 |

Supplemental Figure S2. Onishi et al.

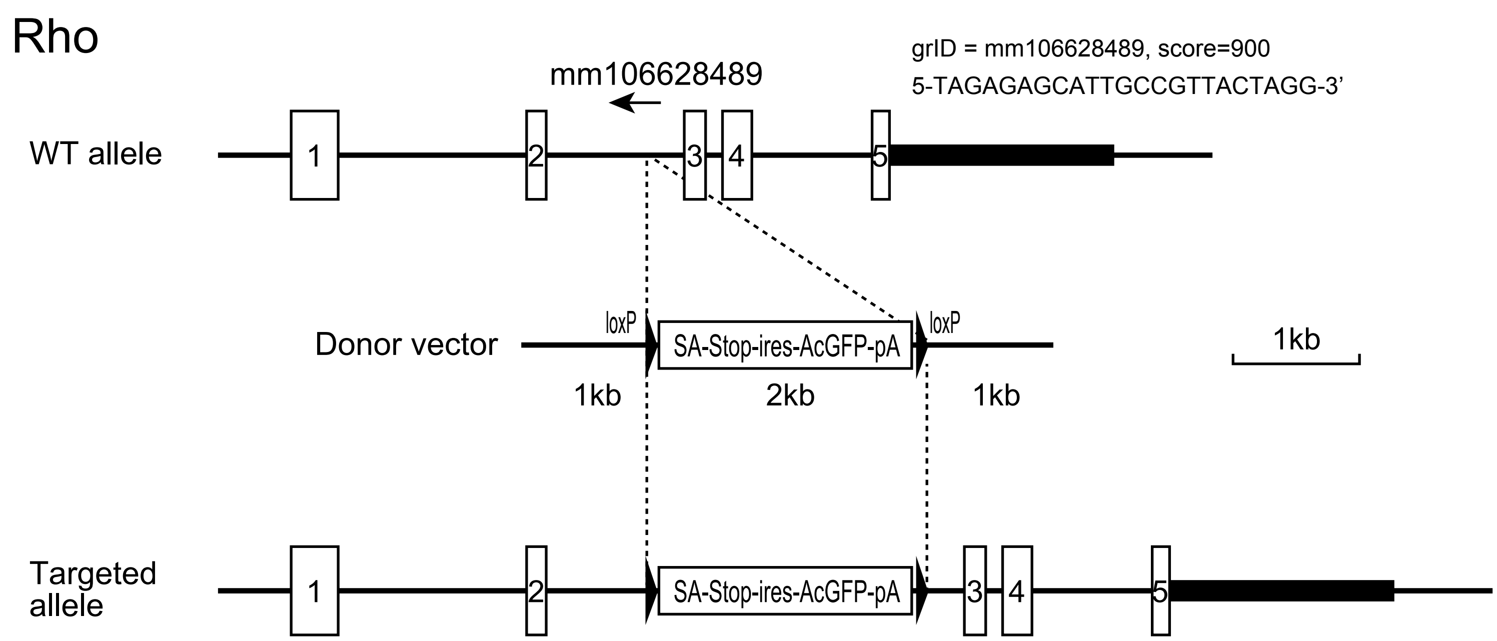


Supplemental Figure S3. Onishi et al.

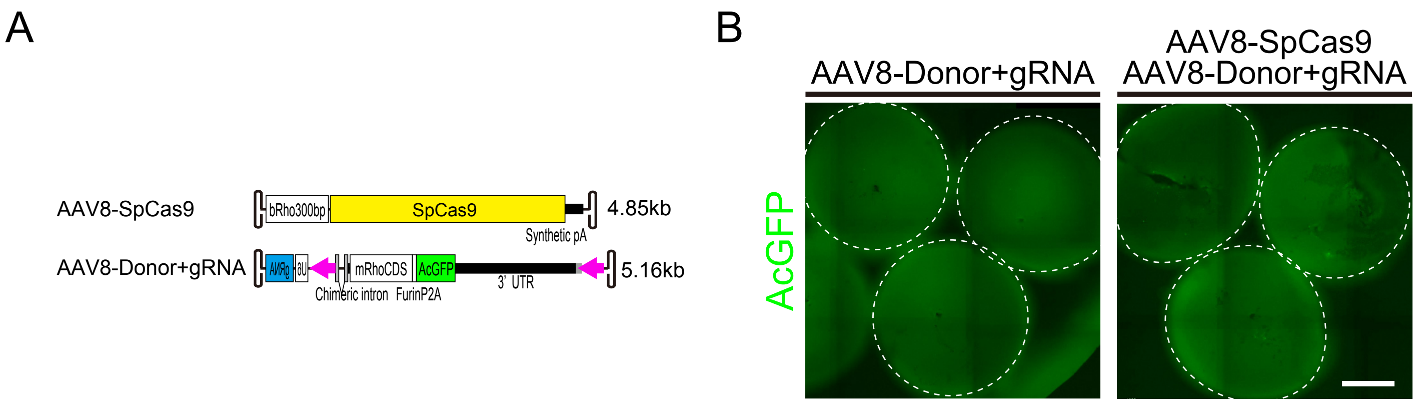


Supplemental Figure S4. Onishi et al.
